# A divergent protein kinase A in the human pathogen *Leishmania* is associated with cell cortex microtubules and controls cell shape

**DOI:** 10.1101/2021.04.24.440790

**Authors:** R. Fischer Weinberger, S. Bachmaier, V. Ober, G.B. Githure, R. Dandugudumula, I.Q. Phan, M. Almoznino, E. Polatoglou, P. Tsigankov, R. Nitzan Koren, P.J. Myler, M. Boshart, D. Zilberstein

## Abstract

Parasitic protozoa of the genus *Leishmania* cycle between the phagolysosome of mammalian macrophages, where they reside as rounded intracellular amastigotes, and the midgut of female sand flies, which they colonize as elongated extracellular promastigotes. Previous studies indicated that protein kinase A (PKA) plays an important role in the initial steps of promastigote development into amastigotes. Here, we describe a novel regulatory subunit of PKA (which we have named PKAR3) that is unique to *Leishmania* and most (but not all) other Kinetoplastea. PKAR3 is localized to subpellicular microtubules (SPMT) in the cell cortex, where it recruits a specific catalytic subunit (PKAC3). Promastigotes of *PKAR3* or *PKAC3* null mutants lose their elongated shape and are round but remain flagellated. Truncation of an N-terminal formin homology-like domain of PKAR3 results in its detachment from the SPMT, also leading to rounded promastigotes. Thus, the tethering of PKAC3 kinase activity via PKAR3 at the cell cortex is essential for maintenance of the elongated shape of promastigotes. This role of PKAR3 is reminiscent of PKARIβ and PKARIIβ binding to microtubules of mammalian neurons, which is essential for the elongation of dendrites and axons, respectively. Interestingly, PKAR3 does not bind cAMP but nucleoside analogs with a very high affinity similar to the PKAR1 isoform of *Trypanosoma*. We propose that these early diverged protists have re-purposed PKA for a novel signaling pathway that spatiotemporally controls microtubule remodeling and cell shape *via* PKA activity.

## Introduction

*Leishmania donovani* is a (trypanosomatid) protozoan parasite that causes kala azar, a fatal form of visceral leishmaniasis in humans (1). These organisms cycle between the midgut of female sand flies, where they reside as flagellated extracellular promastigotes, and the phagolysosomes of mammalian macrophages, where they live as aflagellate amastigotes (2). Infective metacyclic promastigotes are introduced into the host during the sand fly blood meal; after which they are phagocytosed by resident macrophages near the bite site (3,4). Once inside the host phagolysosome, promastigotes encounter two physical cues that distinguish this environment from that of the vector gut: acidic pH (∼5.5) and elevated temperature (∼37°C). Promastigotes process these cues into a signal that initiates development into amastigotes (5–7).

Among the earliest events during *L. donovani* promastigote-to-amastigote differentiation are changes in the phosphorylation profile of many proteins (8). Significantly, while protein kinase A (PKA)-specific phosphorylation is common in promastigotes, most of these phosphoproteins are dephosphorylated within a few minutes after initiation of differentiation into amastigotes (7). This suggests that regulation of PKA activity is important for regulation of *Leishmania* development. PKA-mediated signaling pathways are ubiquitous in eukaryotic cells, being implicated in growth control, development, and metabolism. The canonical PKA regulatory system in vertebrates involves assembly of catalytic (PKAC) and regulatory (PKAR) subunits into an inactive heterotetrameric complex. Dimerization is mediated by a dimerization and docking (D/D) domain in the N-terminal portion of PKAR (9). When cAMP binds the cyclic nucleotide-binding domains (cNBDs) near the C-terminal of each PKAR, the resultant conformational change triggers dissociation of the PKAC/R complex and release of catalytically active PKAC subunits. Compartmentation of PKA activity to different cellular microdomains is achieved (at least in higher eukaryotes) through interaction of the D/D domain of PKAR with a diverse family of A-kinase anchoring proteins (AKAPs) that vary according to biological context (10). However, PKARs from many lower eukaryotes (including trypanosomatids) appear to lack the D/D domain (11,12) and they lack obvious AKAPs (13).

Most eukaryote genomes (with the notable exception of plants and algae) encode one or more PKAC subunits (13). Trypanosomatids (including *L. donovani*) are no exception, having two subunits (PKAC1 and C2) that are almost identical and a third (C3) that is more divergent (14–16). Phosphoproteomic analyses of *L. donovani* revealed that all three PKAC subunits contain at least two phosphorylation sites (17). One, within the kinase active site loop, matches the canonical PKA phosphorylation motif (RXXpS/pT) and is phosphorylated only in promastigotes. The second is located within a canonical ERK1/ERK2 substrate motif (pSP) close to the C-terminus and while PKAC1/C2 is phosphorylated at this site in promastigotes, PKAC3 is phosphorylated only after exposure to the differentiation signal (8). In other eukaryotes, phosphorylation of both a threonine (T_197_) in the activation loop and a serine (S_338_) near the C-terminus is necessary for full activation of PKAC, suggesting that PKAC1/C2 are active in *L. donovani* promastigotes, while PKAC3 is transiently activated only upon receiving the differentiation signal.

Most eukaryotes also possess one or more PKAR subunits, whose number and evolution are more dynamic than for PKAC (13). All trypanosomatids have a regulatory subunit (PKAR1) that has been shown to localize to the flagellum of *Trypanosoma brucei* (18)(19)(20) and *L. donovani*, where it is expressed only in the promastigote stage (7,21). However, in contrast to higher eukaryotes, trypanosomatid PKAR1 appears to be monomeric and unable to bind cAMP (11,12).

Interestingly, *L. donovani* (along with other *Leishmania* species and *Trypanosoma cruzi*, but not *T. brucei*) have a second PKAR-like subunit (14), containing at least 10 sites that are phosphorylated or dephosphorylated during differentiation, implying that it may have a regulatory role (8). Here, we show that this protein (which we have renamed PKAR3) forms a holoenzyme complex with PKAC3 and anchors it to subpellicular microtubules at the cell cortex *via* a formin FH2-like domain at the N-terminus of PKAR3. Promastigotes of *L. donovani* null mutants lacking either PKAR3 or PKAC3 are predominantly rounded compared to wild type (WT) cells, suggesting that tethering of PKA kinase activity to microtubules is necessary for maintenance of elongated cell shape. Thus, this study ties together an evolutionary divergent PKA localized to the microtubule cytoskeleton of *Leishmania* with developmental morphogenesis.

## Results

### Leishmania PKA regulatory subunits are evolutionary divergent

Previous analyses of the PKAR subunits from a wide range of organisms revealed that they have undergone dynamic and divergent evolution during eukaryote history (13). We used BLAST searches to identify PKAR orthologues in 15 different genera and two unclassified trypanosomatids from the Kinetoplastea (**Table S1**). *Leishmania* and most other trypanosomatids contain two paralogues (PKAR1 and PKAR3) that lack the canonical dimerization and docking (D/D) domain found in the N-terminal region of the PKARs of many (but not all) eukaryotes (**Fig. 1A**). Instead, PKAR1 has an RNI-like domain containing Leucine Rich Repeats (LRRs) that may mediate different protein-protein interactions but not homodimerization (11,2). PKAR3 has a long N-terminal region without obvious domain profile signatures. Construction of a phylogenetic tree (**Fig. S1**) revealed that PKAR1 and PKAR3 represent two separate branches distinct from that containing the canonical PKAR(s) found in most other eukaryotes.

**Figure 1:**
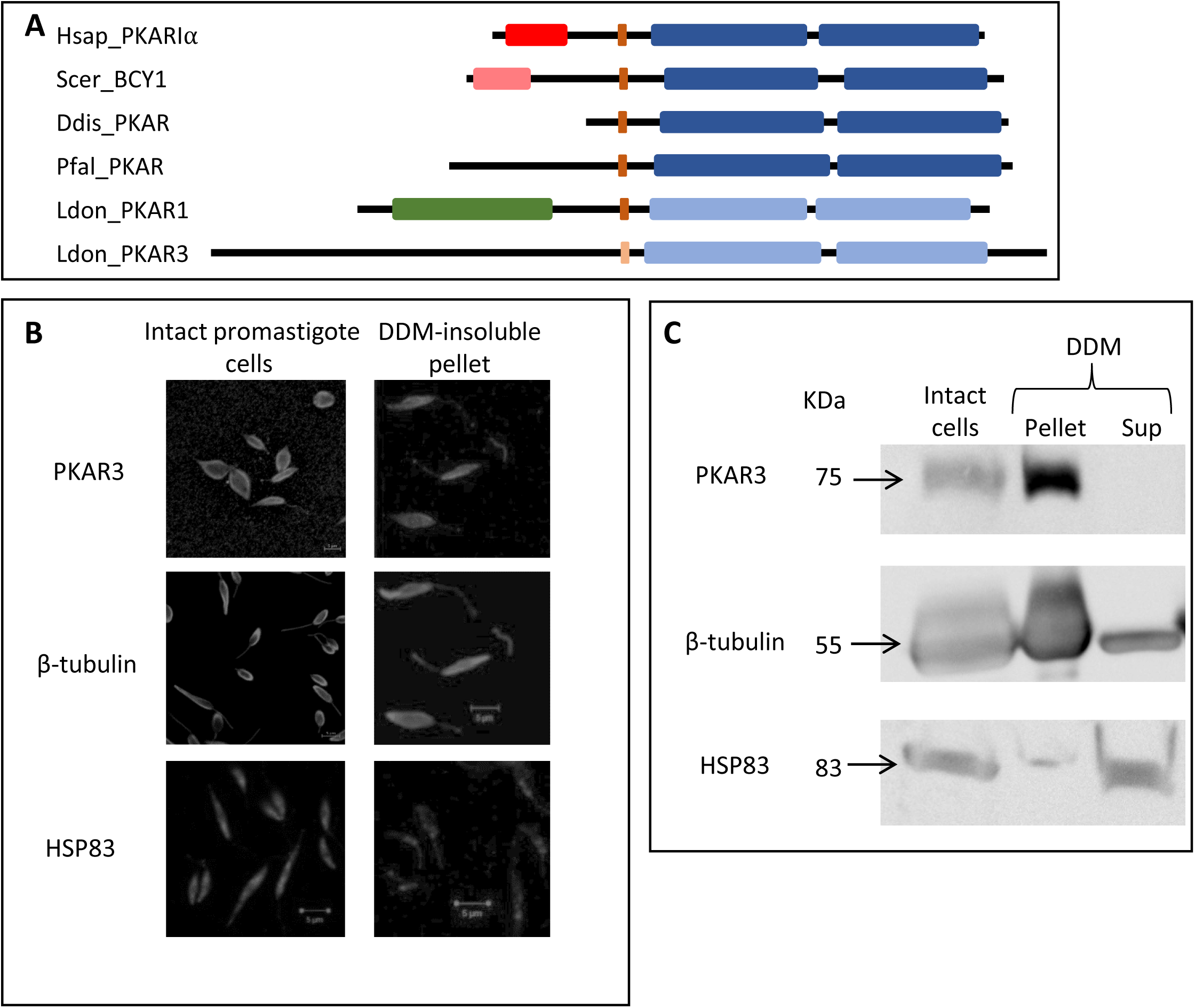
PKAR3 is anchored to the DDM-insoluble subpellicular microtubules (SPMT) at the cortex of *L. donovani* cells. **A**) Schematic representation of selected *Homo sapiens* (Hsap), *Dictyostelium discoideum* (Ddis), *Saccharomyces cerevisiae* (Scer), *Plasmodium falciparum* (Pfal) and *Leishmania donovani* (Ldon) PKAR orthologues showing the dimerization and docking (D/D) domain (red), RNI-like LRR domain (green), (pseudo-)inhibitor site (brown), and nucleoside binding domains (blue). A lighter shade indicates a significant sequence difference from that of the canonical (human) PKAR. **B**) Indirect immunofluorescence of PKAR3 (upper row), β-tubulin (middle row) and HSP83 (bottom row) in intact promastigote cells (left column) or the DDM-insoluble pellet containing the SPMT (right column). Staining was carried out using antibodies against PKAR3, β-tubulin and HSP83. Fluorescence was detected using a Zeiss LSM 700 inverted confocal microscope, with the scale bar representing 5μm. **C)** Proteins from intact or DDM-extracted (supernatant and pellet) promastigotes were separated on 10% SDS-PAGE and the blotted proteins subjected to Western analysis using the antibodies indicated on the left side of the panels.

### Cytoskeletal localization of PKAR3 to cortical subpellicular microtubules

As shown in **Fig. S2**, polyclonal antibodies raised in rabbits against recombinant PKAR3 reacted with a 72 kDa protein in wild-type (WT) cells. This protein was absent in null mutants (Δ*pkar3*), but reappeared after ectopic expression of full-length *PKAR3* in the null mutant (Δ*pkar3::PKAR3_FL_*). Confocal microscopy using the antibody against PKAR3 revealed its localization to the cell cortex in *Leishmania* promastigotes (**Fig. 1B**). The cell cortex contains a rigid cytoskeleton made of stable subpellicular microtubules (SPMT) that are composed largely of β-tubulin (22)(23)(24). Previous studies have shown that non-ionic detergents can solubilize the trypanosomatid surface membrane, while leaving the SPMT intact (25–27). Trématent of *L. donovani* promastigotes with the zwitterionic detergent n-dodecyl β-D-maltoside (DDM, which is neutral at pH 7) resulted in enrichment of β-tubulin in the DDM-insoluble fraction (**Fig. 1B** and **1C**). In contrast, cytosolic HSP83 (28) is absent from the DDM-insoluble fraction. The presence of PKAR3 in the DDM-insoluble pellet suggests that it is associated with the SPMT.

Binding of a PKA regulatory subunit (PKARIβ) to stable microtubules has been documented in neuronal axons of mammalian cells (29), and is mediated by interaction of the dimerization and docking (D/D) domain of PKARIβ with a microtubule associated protein 2 (MAP2) (30). The *Leishmania* genome does not contain an obvious orthologue of MAP2 and PKAR3 lacks the canonical D/D domain, suggesting its localization to the SPTM is mediated by a different mechanism. The N-terminal 50 amino acids of PKAR3 is relatively well-conserved and analysis using HHpred (31,32) revealed structural similarity to three Protein Data Bank (PDB) entries containing a formin homology (FH2) domain (**Fig. S3**). Formins are a diverse family of proteins that participate in polymerization of actin and assembly of microtubule cytoskeletons in many eukaryotes (33), leading us to speculate that this domain might be responsible for binding of PKAR3 to the subpellicular microtubules. To test this hypothesis, we generated null mutants of *L. donovani* expressing an ectopic version of *PKAR3* lacking the N-terminal 90 amino acids (*Δpkar3::PKAR3*_ΔN90_). Western blot analysis confirmed that both the full length and truncated proteins are present in whole cell lysates (**Fig. 2A**). However, only full length PKAR3 remains bound to the DDM-insoluble pellet extracted from these cells, the truncated protein was present only in the DDM supernatant, suggesting that it no longer binds the SPMT. Fluorescence microscopy using antibodies against PKAR3 confirmed that the truncated protein no longer localized to the cortex, being spread throughout the promastigote cell body (**Fig. 2B**). Conversely, when a plasmid containing the N-terminal 90 amino acids of PKAR3 (with a C-terminal HA-tag) was ectopically expressed in the *Δpkar3* mutant *(Δpkar3::PKAR3*_N90_), the fusion protein was enriched in the DDM-insoluble pellet (**Fig. 2C**). Taken together, these results indicate that binding of PKAR3 to the microtubule cytoskeleton is mediated through the N-terminus containing the FH2-like domain.

**Figure 2:**
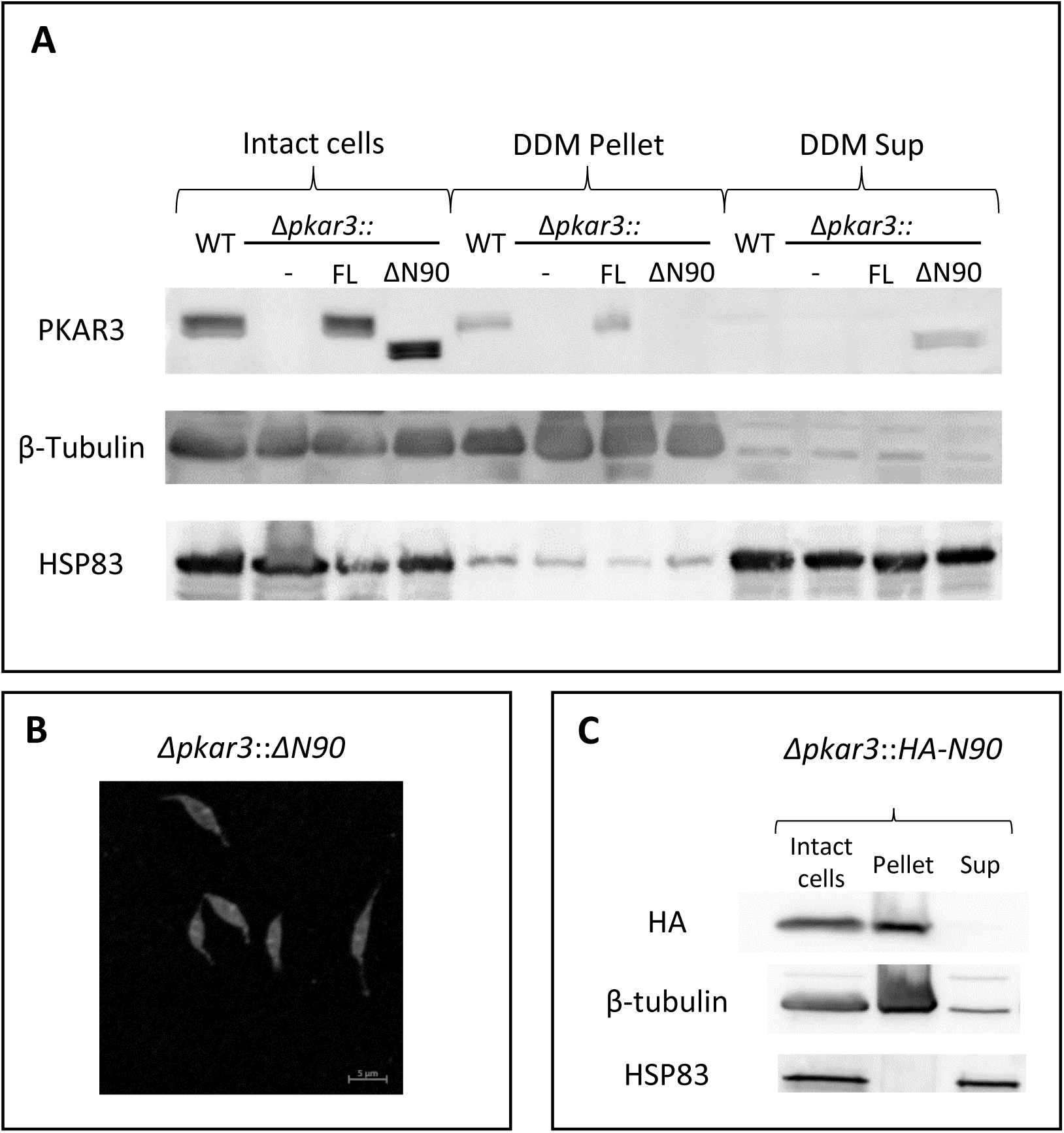
PKAR3 binds to the SPMT *via* an N-terminal FH2-like domain. (**A**) Proteins were extracted from intact and DDM-treated (supernatant and pellet) *L. donovani* promastigotes of the wild type (WT), PKAR3 null mutant (*Δpkar3*), as well as *Δpkar3::FL* and *Δpkar3::ΔN90* cell lines were separated by 10% SDS-PAGE and subsequently subjected to Western blot analysis using anti-PKAR3 (upper row), anti-β tubulin (middle row) and anti-HSP83 (lower row). **B**) Indirect immunofluorescence of PKAR3 in *Δpkar3::ΔN90* promastigotes using antibody against PKAR3. Fluorescence was detected using a Zeiss LSM 700 inverted confocal microscope, with the scale bar representing 5μm. **C)** Proteins were extracted from intact or DDM-treated promastigotes from the *Δpkar3::HA-N90* cell line and subjected to Western blot analysis using antibodies against the HA-tag (upper row), β tubulin (middle row) and HSP83 (lower row).

To further characterize the association of PKAR3 with the SPMT, we used fluorescence resonance energy transfer (FRET) to determine if they are located within close proximity in individual promastigotes (**Fig. 3**). The high value of corrected acceptor FRET intensity observed when PKAR3 and β-tubulin are co-expressed shows that they co-localize within the SPMT (**Fig. 3A**). The low corrected acceptor FRET intensity in the *Δpkar3* mutant validates the approach (**Fig. 3B**). Similarly, we saw low corrected acceptor FRET intensity between β-tubulin and an amino acid transporter (34) located in the cell membrane (**Fig. 3C**). In contrast to the finding above that N-terminal truncated PKAR3 was not retained in the SPMT pellet after DDM treatment, there was significant corrected acceptor FRET detected in *Δpkar3::PKAR3*_ΔN90_ cells (**Fig. 3D**). However, the number of pixels qualified as “interacting” was ∼30% less than obtained with PKAR3_FL_ (**Fig. 3E**). As the concentration of ectopically expressed PKAR3_FL_ and PKAR3_ΔN90_ proteins was high in these cells (**Fig. 2A**), other regions of PKAR3 may also interact with microtubules (albeit more transiently), especially in the absence of detergent. The FRET analyses support our hypothesis that PKAR3 is attached to the SPMT *via* a direct or indirect interaction with β-tubulin.

**Figure 3:**
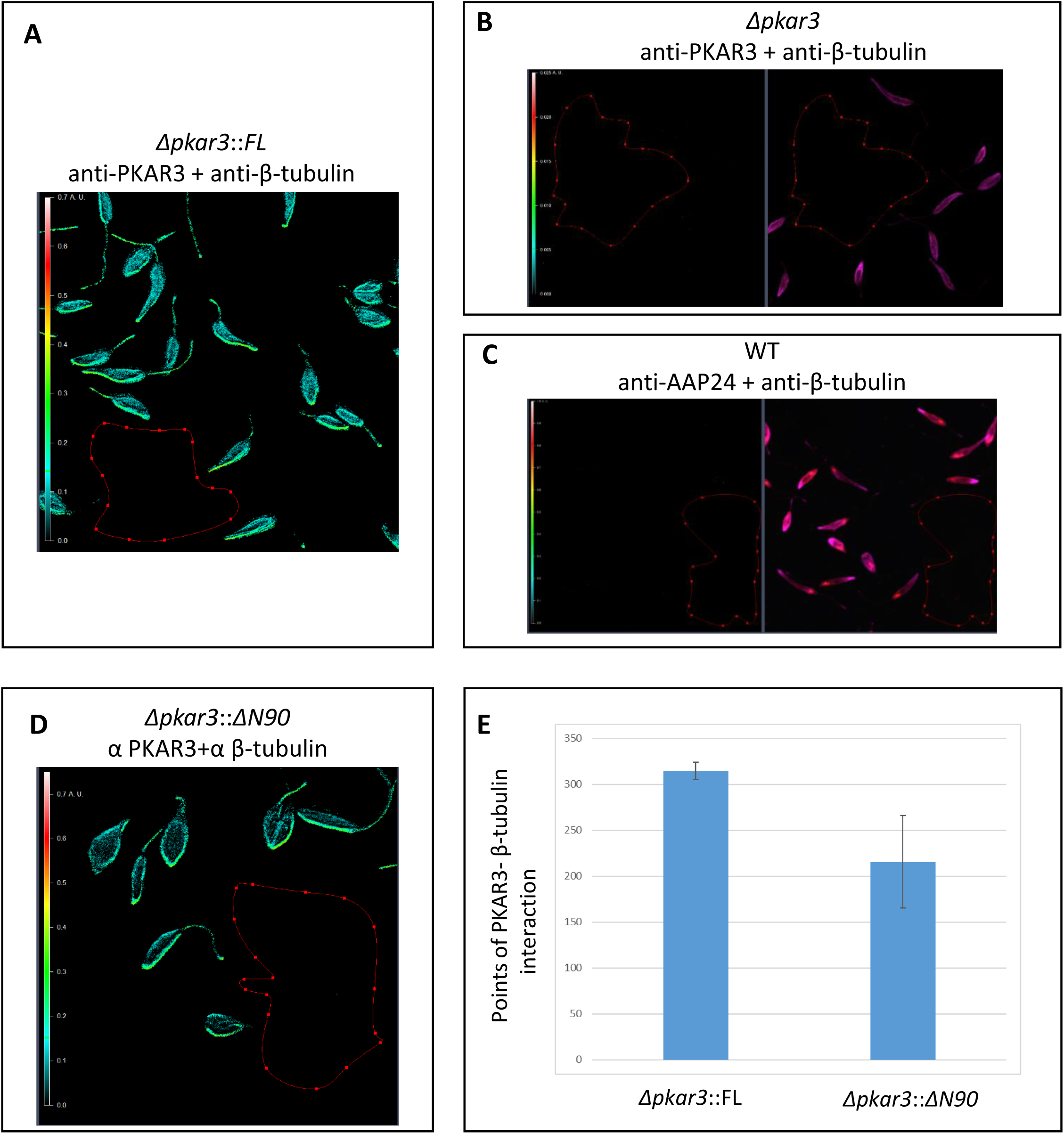
PKAR3 associates with tubulin of the SPMT. **(A)** Promastigotes of *L. donovani* WT were labeled with rabbit anti-PKAR3 and mouse anti-β tubulin, followed by secondary antibody detection with goat anti-rabbit antibodies conjugated to Alexa fluor 568 and goat anti-mouse antibodies conjugated to Alexa fluor 647. Fluorescence Resonance Energy Transfer (FRET) emission was calculated using the Axio Vision program, with the color scale on the left of each panel indicating the strength of FRET emission. **(B)** Null mutants of *PKAR3* (*Δpkar3*) were labeled with the same antibodies as above. FRET analysis is shown in the left panel with cell fluorescence shown in the right panel. (**C)** WT promastigotes were labeled with rabbit antibodies against the AAP24 transporter and mouse antibodies against β-tubulin, followed by secondary antibody detection with goat anti-rabbit antibodies conjugated to Alexa fluor 568 and donkey anti-mouse antibodies conjugated to Alexa fluor 647. FRET analysis is shown in the left panel and fluorescence in the right panel. (**D)** FRET analysis between PKAR3 and β-tubulin in cells expressing *Δpkar3::ΔN90*, carried out as in panels A and B. (**E)** Quantitation of pixels above the interaction threshold FRET value of PKAR3 and β-tubulin as determined from *Δpkar3::FL* and *Δpkar3::ΔN90* add-back cell lines, as described above. One-way ANOVA indicated that number of FRET pixels containing *Δpkar3::ΔN90/*β tubulin were 30% less than *Δpkar3::FL/* β tubulin (P<0.05, n=3)

### PKAR3 associates with PKAC3 to form a holoenzyme complex

Because antibodies against *L. donovani* PKA catalytic submits are not available, we expressed a Ty1-tagged version of *L. donovani* PKAR3 (LdPKAR3-Ty1) under the control of a tetracycline repressor in the *Trypanosoma brucei Δpkar1* cell line (7,12). As expected, expression of LdPKAR3-Ty1 increased in the presence of tetracycline (**Fig. S4A**), with some “leaky” background expression present in the absence of tetracycline. Although the majority of LdPKAR3-Ty1 was found in the DDM-soluble supernatant after detergent fractionation of *T. brucei* bloodstream form cells, the protein was also present in the DDM-insoluble pellet **(Fig. S4B)**, indicating that the association with the SPMT seen in *L. donovani* might be preserved after ectopic expression in *T. brucei*. Affinity purification of LdPKAR3-Ty1 from the DDM supernatant using anti-Ty1 antibody coupled to magnetic beads, followed by Western blotting with antibodies against the PKA catalytic subunits, showed that LdPKAR3 formed a stable complex with TbPKAC3 (**Fig. 4A**), but not with TbPKAC1/2 (**Fig. 4B**). Interestingly, up-regulation of LdPKAR3-Ty1 expression was accompanied by increased TbPKAC3 expression, possibly due to its stabilization in a complex with R3 (**Fig. S4A**). This result was confirmed by co-expression of *L. donovani* PKAR3 with *L. donovani* PKAC3 in the *L. tarentolae* LEXSY expression system (Jena Bioscience, Jena) and purification of the heteromeric complex by tandem affinity chromatography using an N-terminal His_6_-tag on PKAR3 and a Strep-tag on PKAC3 (**Fig. 4C**). As a positive control, we co-purified the *L. donovani* PKAR1*/*C3 complex in a parallel experiment (**Fig. 4D**). We conclude that PKAR3 and PKAC3 form a stable complex similar to the PKAR1/PKAC complexes previously reported in *T. brucei* (12) and validate PKAR3 as subunit of a PKA holoenzyme complex.

**Figure 4:**
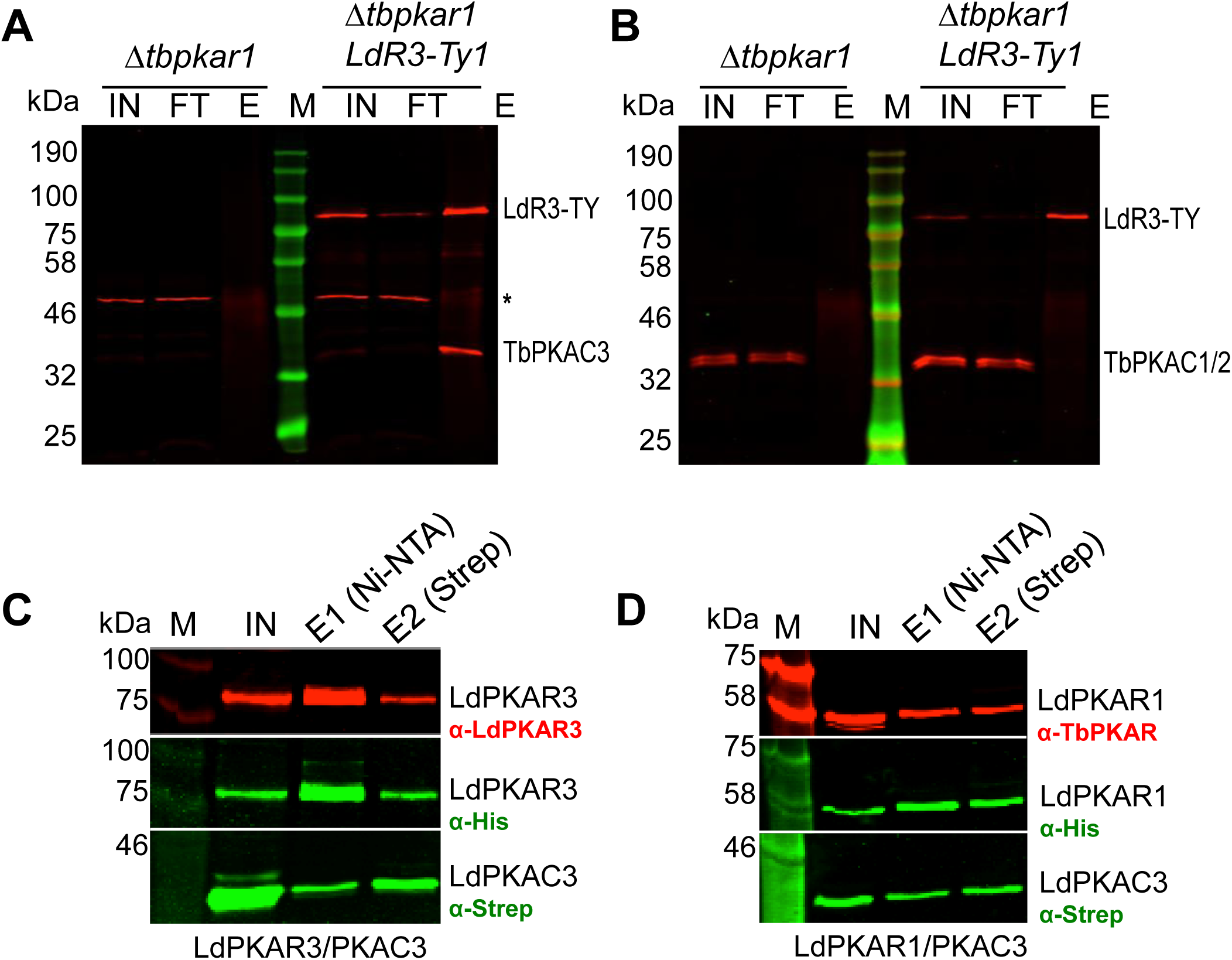
PKAR3 associates with PKAC3. (**A**) Proteins from the DDM-soluble fractions of the *T. brucei* MiTat 1.2 *PKAR1* null mutant (7) parental cell line (*Δtbpkar1*) and those expressing an tetracycline-regulatable version of *L. donovani PKAR3* containing a C-terminal Ty1-tag (*LdR3-Ty1*) were immunoprecipitated with mouse monoclonal antibodies against Ty1, followed by Western blotting with rabbit antibodies against *L. donovani* PKAR3 and *T. brucei* PKAC3. The following abbreviations denote the different fractions interrogated: IN (input), FT (flow-through), and E (elution). The sizes of the molecular weight marker are shown in kilodaltons (kDa). An unidentified protein that cross-reacted with anti-PKAC3 antibody is indicated by an asterisk. (**B**) The same samples were probed with antibodies against PKAR3 and PKAC1. (**C**) *L. donovani* PKAR3 and PKAC3 containing N-terminal His_6_ and Strep tags, respectively, were co-expressed in *L. tarentolae* and soluble proteins affinity purified on tandem Ni-NTA and Strep-Tactin columns. Aliquots from the input (IN) and eluate (E1, E2) fractions were analyzed by Western blotting with rabbit antibodies against LdPKAR3, as well as mouse monoclonal antibodies against the His (Bio-Rad) and Strep (Qiagen) tags. (**D**) A similar experiment performed using tagged *L. donovani* PKAR1 and PKAC3 and probed with rabbit antibodies against TbPKAR1.

To further assess the interaction between PKAR3 and PKAC3 *in vivo*, HA-tagged PKAC3 was ectopically expressed in *L. donovani* WT and *Δpkar3* promastigotes. Western blot analysis of whole cell lysates showed that while PKAC3-HA was expressed at similar levels in both the WT and null mutant; it was detected at high levels in the DDM-insoluble fraction from WT promastigotes, but not in *Δpkar3* (**Fig. 5A**). These results were confirmed by proteomic analysis of the DDM-insoluble fraction of WT and *Δpkar3* promastigotes, which showed that the relative abundance of SPMT-associated PKAC3 was ∼30-fold higher in WT than *Δpkar3* (**Fig. 5B**). In contrast, actin (which is not associated with PKAR3) showed little difference in relative abundance between samples.

**Figure 5:**
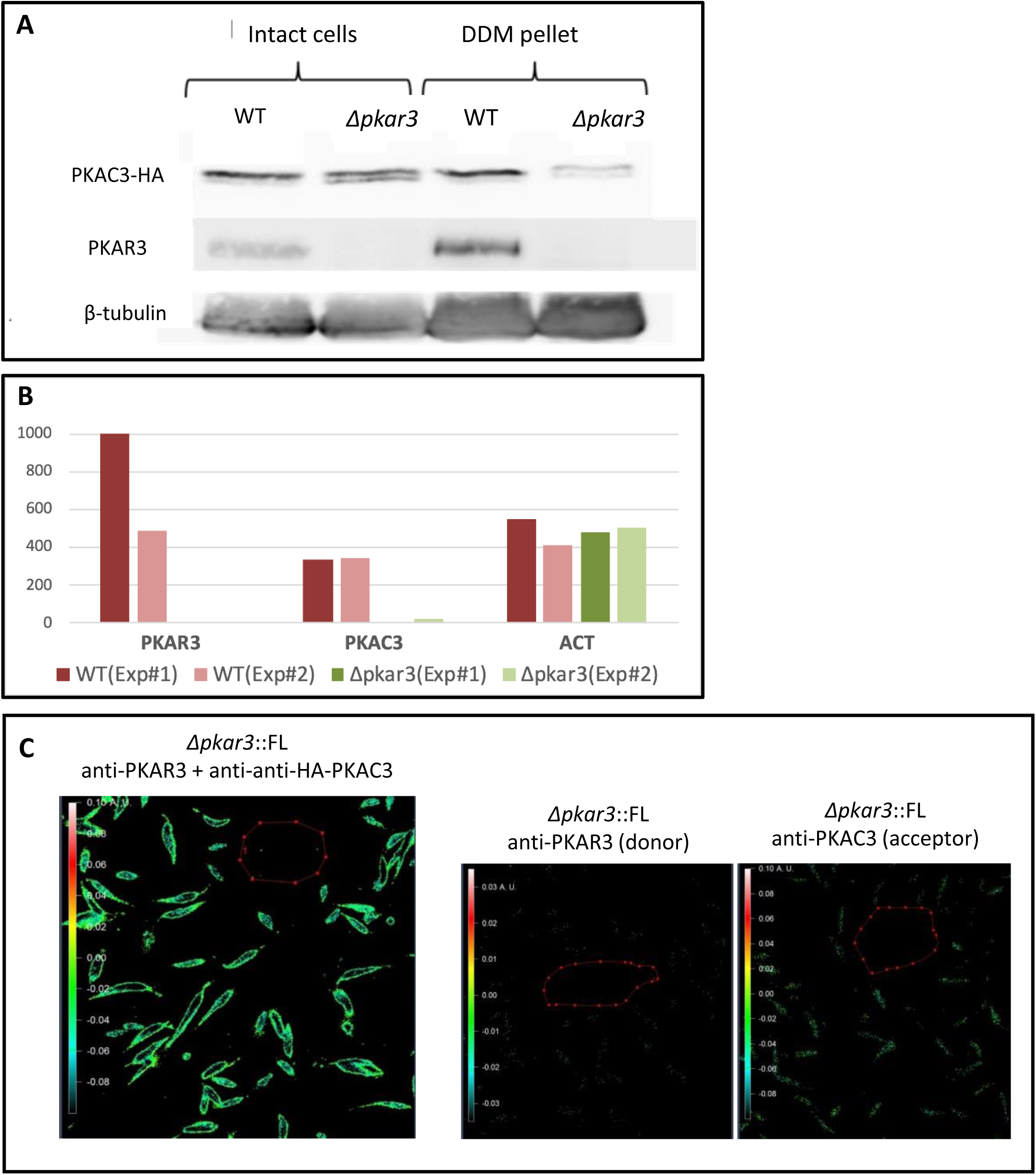
PKAC3 binding to the subpellicular microtubules is mediated by PKAR3. **A)** *L. donovani* promastigote wild type (WT) and PKAR3 null mutants (*Δpkar3*) cell lines expressing HA-tagged PKAC3 were subjected to subpellicular microtubule enrichment using 0.5% DDM, and proteins probed by Western blot using antibodies against the HA-tag (upper row), PKAR3 (middle row) and β-tubulin (lower row). **B**) A gel slice containing proteins with a molecular mass range of 35-42 kDa was excised and subjected to mass-spectroscopy using an Orbitrap (Thermo, Ltd). The relative abundance of peptides from PKAC3, PKAR3, and actin (ACT) are shown for two replicate experiments using WT 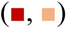 and *Δpkar3* 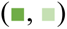 cell lines. **(C)** The WT cell line expressing HA-tagged PKAC3 was labeled with rabbit anti-PKAR3 and mouse anti-HA, followed by goat anti-rabbit antibodies conjugated to Alexa fluor 568 and goat anti-mouse antibodies conjugated to Alexa fluor 647. The left panels show FRET emission color and strength as calculated using the Axio Vision program, with the color scale ruler on the left indicating the strength of interaction between the two labeled proteins. The two right panels show control experiments using antibody against PKAR3 (donor) or HA-tagged PKAC3 (acceptor) alone, followed by the two secondary antibodies together as in the left panel.

We also employed FRET to confirm the molecular proximity of PKAR3 and PKAC3-HA in *L. donovani* promastigotes. As shown in the left panel **of** **Fig. 5C**, the two proteins show a high value of corrected acceptor FRET intensity. Promastigotes labeled with antibodies against the donor (PKAR3) or acceptor (PKAC3-HA) alone, displayed no background FRET (**Fig. 5C**, right panels). These results confirm PKAR3-mediated recruitment of PKAC3 to the SPMT in the parasite cell cortex.

### Structural analysis of the interaction between PKAR3 and PKAC3

The mammalian PKAR and PKAC subunits have been extensively studied at the atomic level, both alone and in complex (35)(36)(37) and an X-ray structure is available for *T. cruzi* PKAR1 (12), allowing us to perform homology modeling based on these structures. Since the N-terminal sequence of PKAR3 is divergent from human (and other) PKAR subunits and is predicted to be largely low-complexity and disordered (except for the N-terminal 50 amino acids), our models were built using only the C-terminal portion including the inhibitor/pseudo-inhibitor sequence and putative nucleoside binding domains. The conserved residues in mammalian cNBDs (R_210_ and R_334_ of PKARIα) have been replaced by N_443_ and A_567_ in PKAR3 (**Fig. 6A** **& B**) and E_560_ in the putative nucleoside binding pocket of latter would likely clash with cAMP phosphate (**Fig. 6B**). We expressed the C-terminal portion of *L. donovani* PKAR3 in *E. coli*, followed by affinity purification and refolding (**Fig. S5**) to produce ligand-free protein for binding assays by isothermal titration calorimetry (ITC) to test whether it binds cAMP. These experiments show a complete absence of cAMP binding under these conditions (**Fig. 6C**) and we found high-affinity binding of the nucleoside analogues 7-cyano-7-deaza-inosine (7CN-7C-Ino, also known as Jaspamycin) and Toyocamycin (Toyo), with dissociation constants of 1 nM and 3.8 nM, respectively (**Fig. 6D**), similar to those for the cAMP-independent PKAR1 of *T. brucei* (12).

**Figure 6:**
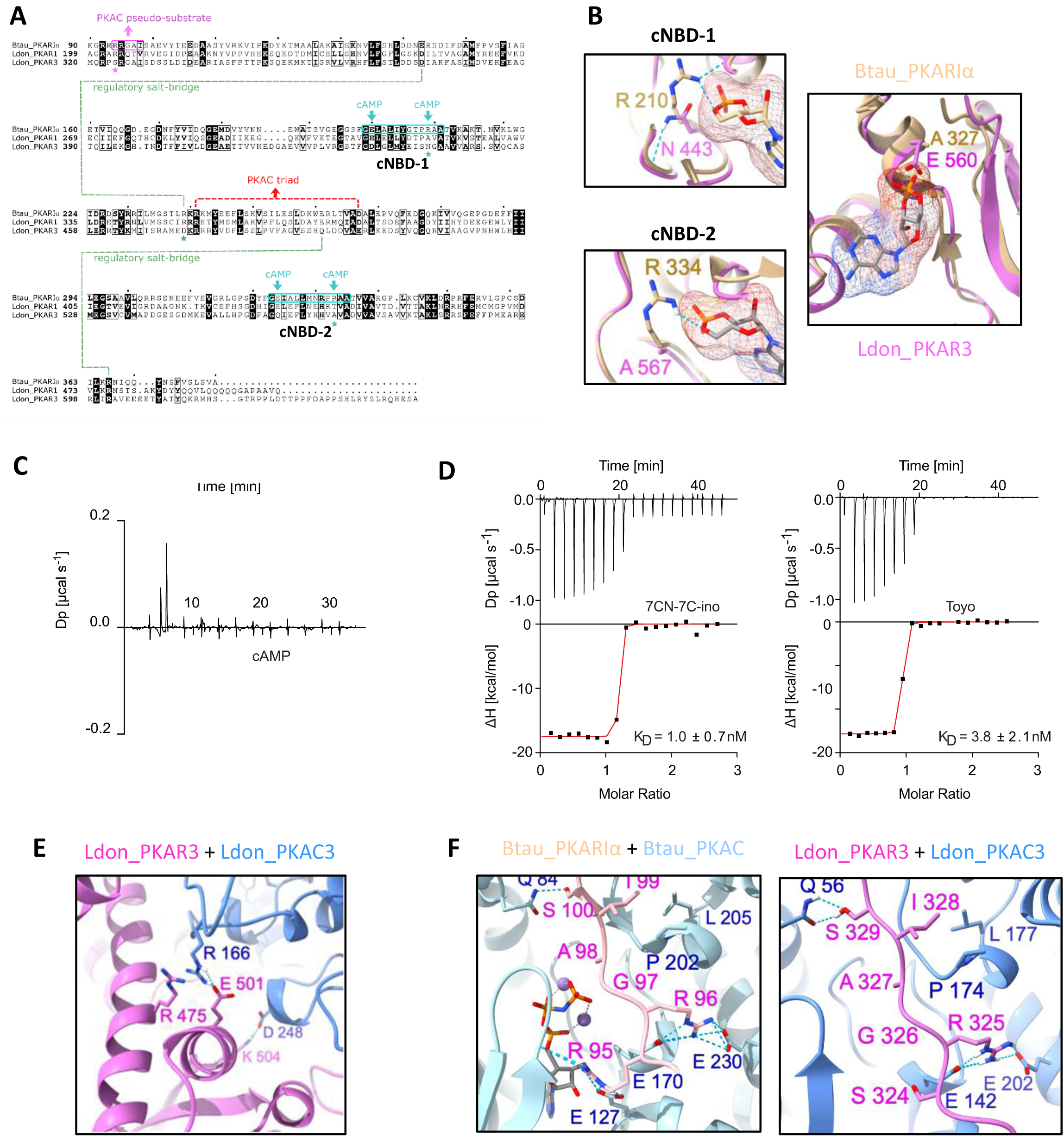
Structural modeling of the PKAR3/C3 complex and ligand binding. (**A**) Protein sequence alignment of *Bos taurus* PKARIα (Btau_PKARIα) with *L. donovani* PKAR1 (Ldon_PKAR1) and PKAR3 (Ldon_PKAR3), indicating regions with known structural importance. Amino acids that are conserved in all three sequences are highlighted in black. (**B**) Structural modelling of the two nucleotide binding domains of Ldon_PKAR3 (pink) based on those from Btau_PKARIα (tan), highlighting the most important changes in the conserved amino acid sequence. (**C**) Binding isotherms from isothermal titration calorimetry (ITC) using refolded ligand-free recombinant protein containing the C-terminal portion (residues 321-647) of *L. donovani* PKAR3 titrated with cAMP. The graph gives the difference power (DP) between the reference and sample cells upon ligand injection as a function of time. The measurement corresponds to noise, excluding binding of cAMP. The results shown are from a representative example of three independent replicates. (**D**) Similar experiments performed with 7CN-7C-inosine (left) or Toyocamycin (right). The upper panels show the difference power (DP) between the reference and sample cells after ligand injection as a function of time, while the lower panels show the total heat exchange per mole of injectant (integrated peak areas from upper panel) plotted against the molar ratio of ligand to protein. The K_D_ is the mean of three or more independent replicates, while the graphs are from a single representative replicate. (**E**) Structural model of the complex formed between Ldon_PKAR3 (pink) and Ldon_PKAC3 (blue) showing amino acids predicted to be involved in the salt-bridges and electrostatic interactions downstream of the pseudo-substrate. (**F**) Side chains of critical residues involved in the PKAR-PKAC interaction for experimentally determined Btau_PKARIα/Cα [PDBID:2QCS] (left) and predicted *Ldon_*PKAR3/C3 (right) complexes.

Electrostatic interaction due to salt-bridges between conserved amino acids have been shown to be important for interaction between PKAR and PKAC subunits in higher eukaryotes (38). The salt-bridge triad between R_240_/D_266_ in bovine PKARIα and R_194_ in PKAC that stabilizes this interaction appears to be conserved in *L. donovani* PKAR3 (R_475_^/^E_501_) and PKAC3 (R_166_) (**Fig. 6E**). However, non-conservative replacement of the first residue (R_95_ in PKARIα to S_324_ in PKAR3) in the inhibitor/pseudo-inhibitor site is predicted to disrupt interaction with negatively charged and polar residues (T_23_, E_99_, E_142_ and Y_302_) in PKAC3 (**Fig. 6F**). Furthermore, changes from E_143_/R_239_ in PKARIα to D_373_/D_473_ in PKAR3 (**Fig. 6A**) likely prevents formation of a salt bridge critical for cAMP-induced allosteric activation of PKARIα (38).

### The PKAR3/C3 complex is necessary for maintaining the elongated shape of promastigotes

Microscopic examination revealed that a significant number of cells in the *PKAR3* null mutant (*Δpkar3*) population lost the normally elongated shape of wild type (WT) promastigotes and became rounded (although still flagellated; **Fig. 7A**). Ectopic expression of full-length (FL), *PKAR3* largely restored the elongated phenotype, but expression of a truncated (ΔN90) version did not. Similarly, *Δpkac3* promastigotes were more rounded than WT, while ectopic expression of full-length *PKAC3* restored the elongated shape. To quantify these changes in shape, we employed Image Streaming flow cytometry (39) to calculate the cell aspect ratio (width over length) of 10,000 cells from each population. ImageStream® analysis identified three different shape groups; elongated (aspect ratio of 0.1-0.4), rounded (aspect ratio of 0.6-1), and ovoid (aspect ratio of 0.4-0.6)(**Fig. S6**). As expected, elongated cells formed the majority in promastigotes (**Fig. S6A,** upper panel), while axenic amastigotes was largely rounded (**Fig. S6A,** lower panel), with the relative numbers of ovoid cells (with an aspect ratio between elongated and round) being similar in both promastigotes and amastigotes. In contrast, promastigotes from the *Δpkar3 and Δpkac3* mutants showed a significantly different distribution of aspect ratio from WT *L. donovani* (**Fig. 7B**). The majority (52%) of WT promastigotes were elongated with 13% rounded; but only 22% of *Δpkar3* promastigotes are elongated, with the proportion of rounded cells increasing to 56%. Ectopic expression of full-length *PKAR3* in the mutant cell-line (*Δpkar3::FL)* partially restored the fraction of elongated cells to 37% and reduced the fraction of rounded cells to 32%. However, mutants expressing the *PKAR3_ΔN90_* add-back were similar to the *Δpkar3* parent, with 46% being rounded and only 17% elongated. *PKAC3* null mutants (*Δpkac3*) also showed a substantial increase in the proportion (53%) of rounded cells with only 14% being elongated. Ectopic expression of full-length *PKAC3* in the *Δpkac3* mutant partially restored the WT phenotype (30% rounded and 31% elongated). These results clearly indicate that both PKAR3 and PKAC3 are critical for maintenance of an elongated cell morphology in *Leishmania* promastigotes and that recruitment of PKAC3 to the SPMT by the N-terminal domain of PKAR3 is critical for maintaining normal morphogenesis in promastigotes.

**Figure 7:**
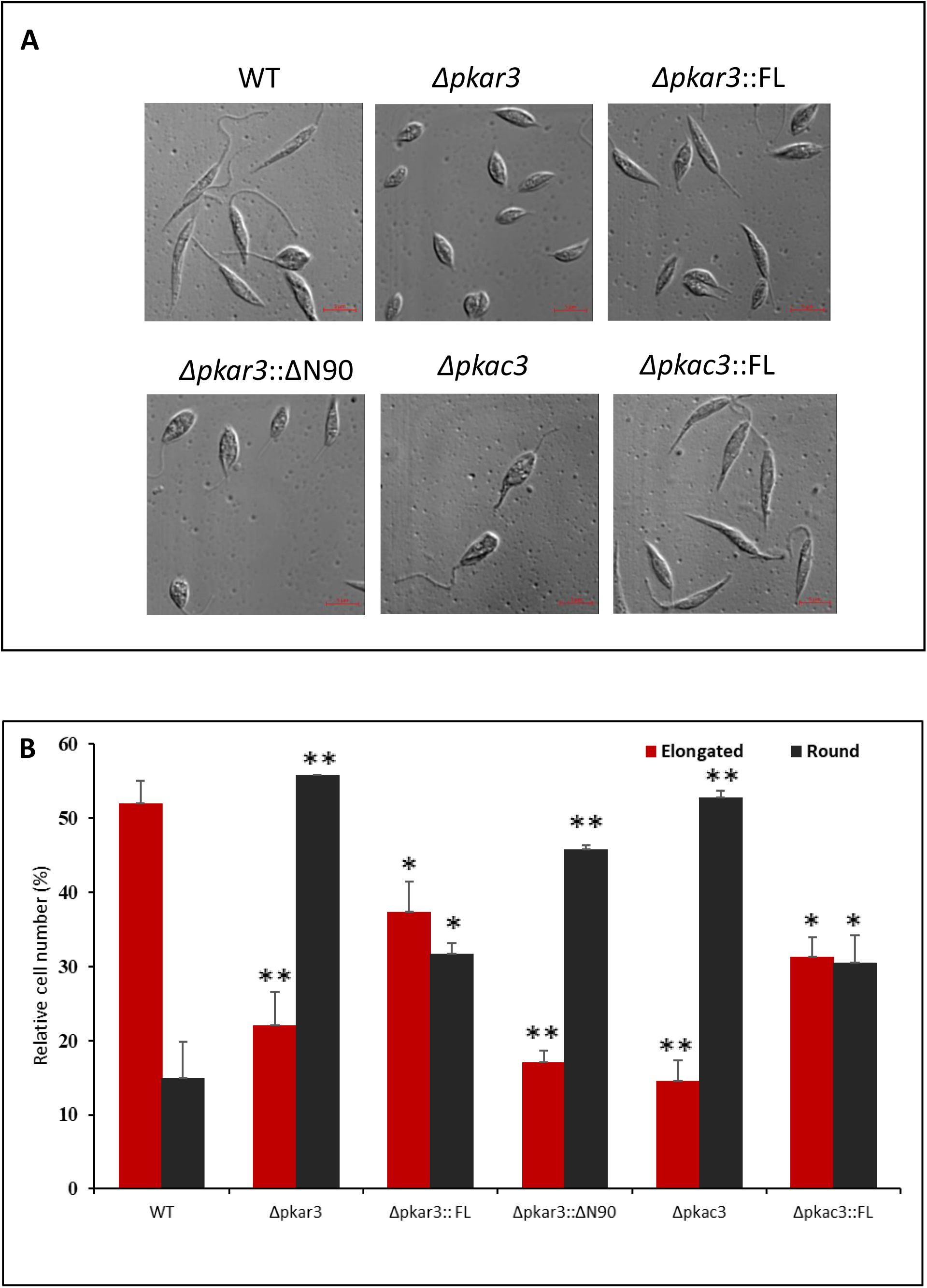
Promastigotes from PKAR3 or PKAC3 mutants are more rounded. (**A**) Microscopic bright field pictures of cells from late-log cultures of *L. donovani* wild type (WT), *PKAR3* null mutant (*Δpkar3)*, *PKAC3* null mutant (*Δpk*ac3), and their add-back derivatives (*Δpkar3::FL*, *Δpkar3::ΔN90*, and *Δpkac3::FL*). **(B**) The percentage of elongated 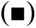 rounded 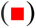 cells in the samples above as determined by image stream analysis. Ovoid cells represented ∼30% in all cell lines and are not included in the graphs. Asterisks indicate statistical significance differences from WT of p <0.05 (*) and p< 0.01 (**), respectively, in three replicate experiments.

## Discussion

In this study, we have functionally characterized a novel regulatory subunit of protein kinase A (PKAR3) that recruits a specific catalytic subunit of PKA (PKAC3) to subpellicular microtubules (SPMT) in the cell cortex of *Leishmania*. This interaction is essential for maintaining the elongated shape of promastigotes, since mutants of *PKAR3* (Δ*pkar3*) become rounded (but remain flagellated), as do those lacking *PKAC3*. Furthermore, truncation of a formin homology (FH2)-like domain at the N-terminal end of PKAR3 resulted in detachment of PKAR3 from the SPMT, again leading to rounded promastigotes. We postulate that the PKAC3 phosphorylates one or more proteins in the cell cortex, allowing precise temporospatial regulation of microtubule remodeling throughout the parasite lifecycle.

In fission yeast, PKA activity triggered by glucose starvation promotes microtubule destabilization by phosphorylation of the microtubule rescue factor CLASP/Cls/Peg1, although this process apparently does not involve recruitment by PKAR (40). Microtubule-associated PKA has also been shown to play a critical role in elongation of axons and dendrites by mammalian neurons (41), with PKARIIβ being responsible for the former and PKARIβ for the latter (30). In this system, microtubule-associated protein 2 (MAP2) binds the D/D domain of the PKARs to link them to the microtubules of neurons (42). However, *Leishmania* lacks an obvious orthologue of MAP2, so it appears that the N-terminal FH2-like domain of PKAR3 compensates by recruiting the PKAR3/C3 complex to microtubules, although whether it binds directly or connects *via* a cryptic anchoring protein remains to be seen.

Homologues of PKAC and PKAR are present in most (but not all) eukaryotes (13), with the canonical PKAR containing an N-terminal D/D domain followed by a linker with the inhibitory sequence and two C-terminal cAMP binding domains. The D/D domain typically dimerizes to form a four-helix bundle that serves as a docking site for different A-kinase-anchoring proteins (AKAPs) that allow the PKAR/C complex to be localized to specific sites in the cell. However, the D/D domain has been lost several times during eukaryotic evolution, including in trypanosomatids, where PKAR1 contains an RNI-like LRR domain and PKAR3 contains a long, mostly disordered, N-terminal region. Interestingly, *Paratrypanosoma confusum* (a trypanosomatid parasite isolated from mosquitoes) has an additional paralogue of PKAR (see **Table S1**) that is more closely related to those found in other eukaryotes (**Fig. S1**). All three paralogues are also found in *Bodo saltans* (a free-living kinetoplastid), suggesting that PKAR1 and PKAR3 arose by gene duplication (and subsequent domain shuffling) early in the evolution of the Kinetoplastea. Interestingly, PKAR3 has been lost in several trypanosomatid genera; *Phytomonas* and *Vickermania* (monoxenous insect and/or plant pathogens), as well as four sub-genera (*Duttonella*, *Herpetosoma*, *Nannomonas* and *Trypanozoon*) of *Trypanosoma (*extra-cellular parasites of mammals). Since many of these species appear to lack amastigotes, this pattern is consistent with the hypothesis that PKAR3 plays an important role in regulating changes in cell morphology during intracellular development.

It has been previously shown that sequence changes in the cNBDs of PKAR1 make it refractory to cAMP binding (12). Here we show that PKAR3 is equally refractory to cAMP and that two high-affinity ligands of PKAR1 (7-CN-7C-ino and toyocamycin) also bind PKAR3 with affinities in the nanomolar range. However, attempts to dissociate the PKAR3-PKAC3 complex using 7-CN-7-C-ino treatment were unsuccessful, leaving it open whether PKAR3 is a ligand-controlled inhibitor of PKAC3. Indeed, the first loop of the pseudo-inhibitor domain of PKAR3 contains a serine instead of the arginine conserved in other PKARs, suggesting that it cannot dock the PKAC3 active site. PKAC3 may remain catalytically active while complexed with PKAR3, allowing maintenance of the elongated shape of promastigotes. Active PKAR/C complexes have also been shown in higher mammals (30). It is possible that *Leishmania* (and other Kinetoplastidae) utilize PKAR3 primarily as an anchoring or tethering subunit to localize a specific PKA catalytic subunit close to its targets in the cell. In a similar way PKAR1 seems to tether PKAC1 and 2 to the flagellum in *Leishmania* promastigotes and in *Trypanosoma* (7,18-20). Whether the nucleoside ligands of PKAR1 and PKAR3 regulate kinase activity of PKAC1/2 and PKAC3, respectively, in the functional context of an upstream pathway remains to be determined. Alternatively, the temporal control of kinase activity might operate via the catalytic subunit and the role of PKAR3 be limited to the tethering function, i.e. the spatial control. This is indeed an attractive speculation as it has been shown that PKAC3 mRNA (previously named c-lpk2) is downregulated upon induced promastigote to amastigote differentiation with very fast kinetics (15). This model is perfectly compatible with the observed rounding upon induction of differentiation and the mutant phenotypes.

In conclusion, we have shown that *Leishmania* (and likely other trypanosomatid parasites) have re-purposed PKA for a cAMP-independent mechanism to spatiotemporally regulate microtubule remodeling and cell shape during parasite development upon host infection.

## Author contributions

RFW, SB, RD, MA, VO, GBG, EP, PT and RNK conducted experiments. IQP conducted structure modeling. PJM supervised these efforts and performed the phylogenetic analyses. MB supervised the work in *T. brucei*, *L. tarentolae* and the ITC experiments. DZ supervised the experiments in *L. donovani* and managed the overall project. He also wrote the manuscript together with MB, PJM and SB.

## STAR Methods

### Phylogenetic analysis of PKA regulatory subunits

The amino acid sequences of PKAR orthologues were obtained from 19 kinetoplastid genomes representing 17 different genera and two unclassified Trypanosomatidae (**Supplementary Table S1**), as well as 8 other eukaryotic species (**Supplementary Table S2**). In several cases, tBlastn searches of unannotated genomes were performed to identify the open reading frame(s) encoding PKAR proteins. Protein domains within each sequence were identified using the InterProScan (43) module of Geneious Prime (https://www.geneious.com) and/or the HHpred website of the MPI Bioinformatics Toolkit (31,32). Protein sequences were aligned using the Clustal Omega (44) of Geneious and phylogenetic trees constructed using the RAxML module implemented in Rapid Bootstrapping mode with 100 trees using the GAMMA BLOSUM62 substitution matrix (45).

### Cell culture

*L. donovani* MHOM/SD/00/1S (28) axenic promastigotes were grown in Earle’s-based medium 199 (M199; Biological Industries, Ltd) supplemented with 10% heat-inactivated Fetal Bovine Serum (GIBCO, Ltd) and 1% Penicillin-Streptomycin solution (Biological Industries, Ltd). Trypanosomes were cultivated at 37°C and 5% CO_2_ in modified HMI-9 culture medium (46) supplemented with 10% heat-inactivated fetal bovine serum. Cell density was kept below 1×10^6^ cells/ml by regular dilution.

### Detergent enrichment of subpellicular microtubules

Axenic *L. donovani* promastigote (20 ml of cells at 1×10^7^ cells/ml) were washed twice with ice-cold phosphate buffered saline (PBS). The final pellet was suspended to 200 µl PBS containing protease inhibitors mix (Roche cOmplete protease inhibitor) and 0.5% n-dodecyl β-D-maltoside (DDM) in Eppendorf tubes. After a short vortex, the tubes were placed on ice for 15 min, and subsequently centrifuged. The pellet was washed twice with 1 ml of ice-cold PBS containing the protease inhibitors. For immunofluorescence analysis, the pellet was suspended in 100 µl PBS. For Western blot analysis, the pellet was suspended in 100 µl Laemmli buffer. *T. brucei* subpellicular microtubules were enriched employing the same protocol used for *Leishmania*.

### Generation of a polyclonal anti-PKAR3 antibody

The full-length CDS of R3 (encoded by *LinJ.34.2680*) was PCR amplified from genomic DNA of *L. infantum* strain JPCM5 using primers (**Table S2**) designed to introduce an N-terminal His_6_ tag followed by a TEV protease cleavage site and cloned *via Bam*HI and *Not*I restriction sites into pETDuet-1 (Novagen, Merck Millipore). The protein was expressed in *E. coli* Rosetta and purified using Ni-NTA columns followed by TEV protease cleavage. Recombinant R3 was injected to two rabbits with 5 mg/ml protein each. Rabbits were boosted four times with 5 mg/ml of the recombinant R3. Injections and bleedings were carried out as a contracted service (Sigma-Aldrich, Ltd).

### Western blotting

Western blot analysis was done as described previously (47), using a 1:1,000 dilution of polyclonal rabbit anti-R3, anti-TbPKAR1 (7), anti-TbPKAC1/C2 (12) and anti-TbSAXO (25). Rabbit antibodies against TbPKAC3 were used at 1:250 dilution (12). For Western blot detection of proteins recombinantly expressed in *L. tarentolae*, mouse anti-His (Bio-Rad) and mouse anti-Strep (Qiagen) were used. The secondary antibodies IRDye680LT goat anti-rabbit (1:50000; LICOR) and IRDye800CW goat anti-mouse (1:10000; LICOR) were used for detection with the Odyssey^TM^ CLx imaging system (LICOR). Rabbit antiserum against *L. donovani* HSP83 (28) or β-tubulin (cell signalling, Ltd) were used as protein loading marker.

### Gene knockouts

The *PKAR3* gene was deleted from the 1S-2D clone of *L. donovani* using homologous recombination as described previously by Inbar et al. (34). PCR analyses of genomic DNA was used to confirm the absence of the targeted gene (**Fig. S1A**). Polyclonal antibodies raised in rabbits against recombinant PKAR3 reacted with a protein of the correct molecular mass (72 kDa) in Western blots of wild type cells, but not *Δpkar3* mutants (**Fig. S1B**). The protein was present (at slightly elevated levels) in add-back mutants (*Δpkar3::*FL) ectopically expressing full-length R3. The *PKAC3* gene was deleted using the same strategy as above. PCR analyses of *L. donovani* genomic DNA extracted from single colonies confirmed that both copies of the gene are missing from *Δpkac3*, while the antibiotic resistance genes that were inserted to replace *PKAC3* are in place (**Fig. S7**).

### Immunofluorescence

Indirect immunofluorescence analysis was carried out following the methods described by Inbar et al. (34). Briefly, late log phase promastigotes were washed twice in phosphate buffered saline (PBS) and then fixed in 1% formaldehyde/PBS on a slide for 10 min before permeabilization by exposure to 0.2% TritonX-100/PBS for 10 min. Cells were incubated with blocking solution [5% (v/v) non-fat dried skimmed milk powder/PBST] for 30 min at room temperature, incubated with rabbit anti-PKAR3 (this study), rabbit anti-promastigote membrane proteins (28) or mice anti-HA tag antibodies (1:500 dilution) for one hour. Subsequently, slides were washed three times with PBS-Tween and incubated with the fluorescent secondary antibody in darkness for 30 min. Slides were washed three time with PBS-Tween and a drop of DAPI Fluoromount G (Southern Biotech Ltd) was added. Slides were covered with slips, sealed, and then kept in darkness. Subsequently, they were examined using a Zeiss LSM 700 inverted confocal laser scanning microscope. Image processing was done using Zen lite software, Zeiss.

### Fluorescence Resonance Energy Transfer (FRET)

Late log phase *L. donovani* promastigotes ectopically expressing PKAR3 and PKAC3-HA were fixed in PBS containing 4% paraformaldehyde and then settled on slides. PKAR3 was labelled using polyclonal anti-PKAR3 antibody (1:500), followed by detection with goat anti-rabbit Alexa 568 (1:500). PKAC3 was labelled with anti-HA antibody (1:1000, BioLegend, Ltd) and secondary goat anti-mouse Alexa 647 (1:500). Confocal microscopy was obtained using a Zeiss LSM 700 Inverted confocal laser scanning microscope. An excitation wavelength of 578 nm and an emission wavelength of 640 nm and below were used for Alexa568, whereas an excitation wavelength of 651 nm and an emission wavelength of 640 nm and above were used for Alexa647. FRET was assessed with AxioVision FRET software, using the mathematical approach - Youvan’s technique. Youvan’s technique (Fc) is the basis of all correction measurement techniques (out of 4 techniques) in AxioVision FRET. It refers to corrected FRET values when three FRET filter sets are used. The measured values are corrected for the background (bg) and for the crosstalk from the donor (don) and the acceptor (acc). Calculation of the intensity values for FRET follows the formula: Fc = (fretgv - bg) - cfdon (dongv - bg) - cfacc (accgv - bg), where gv is the fluorescence value in every pixel and cf is a constant, which is calculated for each of the samples with only one secondary antibody. Subsequently, the number of interacting molecules in the FRET images were counted using the microscopic analysis software, IMARIS version 9.3.

### Image streaming flow cytometry

Analysis was carried out using Amnis® ImageStream®X Mk II)(48)(49)(50). The Image Streamer was calibrated by feeding it with late log phase axenic promastigotes of wild type and mutants. Late log phase *L. donovani* promastigotes were washed once using ice cold phosphate buffered saline (PBS). Subsequently, cells were fixed by suspending them in PBS containing 4% paraformaldehyde. An aliquot of these cell suspension was injected into the Imagestream Mark II. Each cell was subjected to a snapshot in bright field. The data was then analyzed using IDEAS software, version 6.10, with compensation according to the software standards. Cells were gated for single cells in focus. Further, cells were classified based on shape (elongated, ovoid, round) by considering aspect ratio vs symmetry features. Aspect ratio feature assesses the elongatedness of cells by height/width ratio. Symmetry 2 measures the tendency of the object to have single axis of elongation.

### Inducible overexpression of PKAR3 in Trypanosoma brucei

The *PKAR3* CDS was PCR amplified from genomic DNA of *L. donovani* promastigotes using primers R3_HindIII_fw and R3_lr_Ty1_BamHI_rev (**Table S2**) to introduce a C-terminal Ty1 tag. The PCR product was cloned into the *Hin*dIII and *Bam*HI sites of pHD615 (51), which was then linearized with *Not*I for transfection of a previously described homozygous *T. brucei PKAR1* deletion mutant (7) that additionally expresses a double tetracycline repressor (pHD1313; (52). Transfected cells were selected with 2 µg/ml blasticidin (pHD615) and 2.5 µg/ml phleomycin (pHD1313). PKAR3 expression was induced with 1 µg/ml tetracycline.

### Immunoprecipitation in T. brucei

Immunoprecipitation of PKAR3-Ty1 was carried out by binding anti-Ty1 (53) to magnetic protein A beads (Dynabeads, Invitrogen) followed by a 2-hour incubation with 1×10^8^ trypanosomes lysed in lysis buffer (10 mM Tris/Cl pH 7.5; 150 mM NaCl; 0.5 mM EDTA; 0.5% NP-40; Roche cOmplete protease inhibitor) for 30 min at 4°C. Beads were washed 4x with lysis buffer and proteins were eluted by incubation with 50 µl 2× Laemmli sample buffer for 5 min at 95°C.

### Tandem affinity purification from Leishmania tarentolae

PKAR1 and *PKA*R3 were N-terminally tagged with a hexa-histidine peptide by PCR amplification from *L. donovani* genomic DNA. The PCR products were inserted in the pLEXSY_I-ble3^®^ vector (Jena Bioscience). PKAC3 was N-terminally strep-tagged and inserted into pLEXSY_I-neo3^®^. The vectors were then linearized with *Swa*I and stably co-transfected for holoenzyme expression (PKAR1 and PKAC3 or PKAR3 and PKAC3) into the LEXSY T7-TR^®^ cell line (Jena Bioscience). Protein co-expression, parasite cultivation and protein purification were performed as described in (12) with a few modifications. Briefly, co-expression was induced with 10 µg/ml tetracycline for 48 hours. The cells were harvested by centrifugation (2000×g for 5 min), washed with PBS, and lysed in His binding buffer (50 mM NaH_2_PO_4_ pH 8, 300 mM NaCl, 10 mM imidazole, 1% Triton-X, protease inhibitor cocktail). The soluble fraction was loaded on to a gravity flow Ni-NTA column (Thermo Fisher Scientific). The column was washed twice with His wash buffer (50 mM NaH_2_PO_4_ pH 8, 300 mM NaCl, 20 mM imidazole) prior to elution of the kinase complex with His elution buffer (50 mM NaH_2_PO_4_ pH 8, 300 mM NaCl, 250 mM imidazole). The Ni-NTA eluate was loaded on to a gravity flow Strep-Tactin column (IBA), which was then washed with Strep wash buffer (50 mM NaH_2_PO_4_ pH 8, 300 mM NaCl). Lastly, the kinase complexes were eluted with Strep elution buffer (50 mM NaH_2_PO_4_ pH 8, 150 mM NaCl, 50 mM biotin). The integrity of the co-expressed holoenzyme complexes was analysed by western blot.

### Pulldown of subpellicular microtubules-bound proteins (proteomics)

These experiments aimed to determine the levels of PKAC3 binding to the subpellicular microtubules of *L. donovani* wild type (WT) and *Δpkar3*. Microtubules of WT and *Δpkar3* promastigotes were DDM-enriched as described above. Subsequently, proteins associated with this fraction were separated on 9% SDS-PAGE and a slice containing proteins at the molecular mass range of 35-42 kDa was excised and subsequently subjected to mass spectrometry. The proteins in the gel were reduced with 3 mM DTT (60°C for 30 min), modified with 10 mM iodoacetamide in 100 mM ammonium bicarbonate (in the dark, room temperature for 30 min) and digested in 10% acetonitrile and 10 mM ammonium bicarbonate with modified trypsin (Promega) at a 1:10 enzyme-to-substrate ratio, overnight at 37°C. The resulted peptides were desalted using C18 tips (Homemade stage tips) dried and resuspended in 0.1% formic acid. The peptides were resolved by reverse-phase chromatography on 0.075 X 180-mm fused silica capillaries (J&W) packed with Reprosil reversed phase material (Dr Maisch GmbH, Germany). The peptides were eluted with linear 60 minutes gradient of 5 to 28%, 15 minutes gradient of 28 to 95% and 15 minutes at 95% acetonitrile with 0.1% formic acid in water at flow rates of 0.15 μl/min. Mass spectrometry was performed by Q Exactive plus mass spectrometer (Thermo) in a positive mode using repetitively full MS scan followed by collision induced dissociation (HCD) of the 10 most dominant ions selected from the first MS scan. The mass spectrometry data was analyzed using Proteome Discoverer 1.4 software with Sequest (Thermo) algorithm against *L. donovani* and *L. infantum* proteomes from TriTryp database with mass tolerance of 20 ppm for the precursor masses and 0.05 Da for the fragment ions (TriTrypDB). Oxidation on Met, and Phosphorylation on Ser, Thr, Tyr were accepted as variable modifications and Carbamidomethyl on Cys was accepted as static modifications. Minimal peptide length was set to six amino acids and a maximum of two miscleavages was allowed. Peptide- and protein-level false discovery rates (FDRs) were filtered to 1% using the target-decoy strategy. Semi-quantitation was done by calculating the peak area of each peptide based on its extracted ion currents (XICs) and the area of the protein that is the average of the three most intense peptides from each protein. The mass spectrometry proteomics data have been deposited to the ProteomeXchange Consortium via the PRIDE (54) partner repository with the dataset identifier PXD025222.

### Molecular modeling

PKAR3 homology models were built from each of the PDB templates 2QCS, 5JR7 and 4MX3 with Chimera Modeller (55,56). Multiple sequence alignments were produced with Clustal Omega (44). The PKAR3/C3 complex was predicted by AlphaFold2 using MMseqs2 on ColabFold v1.5.2, with pdb70 templates, alphafold2_multimer_v3 weights for complex predictions using 20 recycles and tolerance set to 0.5, followed by AMBER relaxation (57). Structure images were produced with UCSF Chimera (58).

### Ligand binding studies by isothermal titration calorimetry (ITC)

Protein purification was performed as described by (12) with the following modification: N-terminally truncated LdPKAR3 (aa 321-647) was cloned into pET11_Sumo3 (NEB) with an N-terminal Sumo3-tag and expressed in *E. coli* Rosetta (DE3). Purification of R3(321–647) by Ni-NTA affinity chromatography was followed by SenP2 protease mediated cleavage of the N-terminal Sumo3-tag during dialysis of the protein in 50mM HEPES pH 7.5 and 50mM NaCl. The cleaved protein was either stored at -80°C or directly denatured and separated from prebound ligands as described (12). For ligand binding studies by ITC the protein was further dialyzed against 50mM Tris-Cl pH 8.5, 9.6mM NaCl, 0.4mM KCl, 2mM MgCl2, 2mM CaCl2, 0.5M Arginine, 0.75M Guanidine-HCl, 0.4M Sucrose, 10mM DTT over night at 4°C and subsequently purified by gel filtration chromatography on a Superdex Increase 10/300 GL column (GE healthcare). Refolded and purified proteins were diluted to 10-20 µM in ITC buffer (50mM HEPES pH 7.5 50mM NaCl and 1% DMSO). 100-200 µM 7-cyano-7-deaza-inosine (7CN-7C-ino) or Toyocamycin (Toyo) was diluted in the same buffer. ITC measurements were carried out on a MicroCal PEAQ-ITC (Malvern instrument). 2-4 µl of ligand were injected in a series of 13-19 injections into the protein sample at 298K. The Differential Power (DP) between reference and sample cell was maintained at 8-10 µcal s–^1^ in all experiments. Data analysis was performed with the MicroCal PEAQ-ITC software applying a model with one binding site.

## Acknowledgements

We thank the Smoler Proteomics Center at the Technion for all proteomic analyses. Funding for this work was provided by the USA-Israel Binational Science Foundation (BSF), grant number 2017030 to DZ and PJM, the German-Israel Foundation (GIF) grant number I-112-416.10-2017 to DZ and MB, contract HHSN272201700059C from the National Institute of Allergy and Infectious Diseases to PJM, and grant 16GW0281K from the Federal Ministry of Education and Science of Germany to MB.

