## Supplementary figures for "A divergent protein kinase A in the human pathogen *Leishmania* is associated with cell cortex microtubules and controls cell shape"

Figure S1

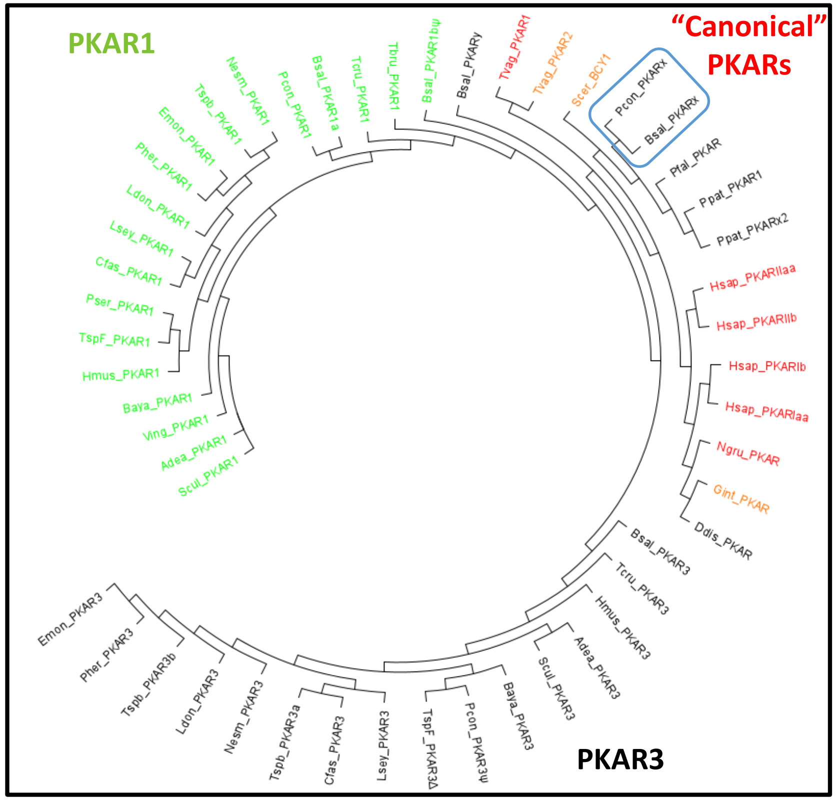

Figure S2

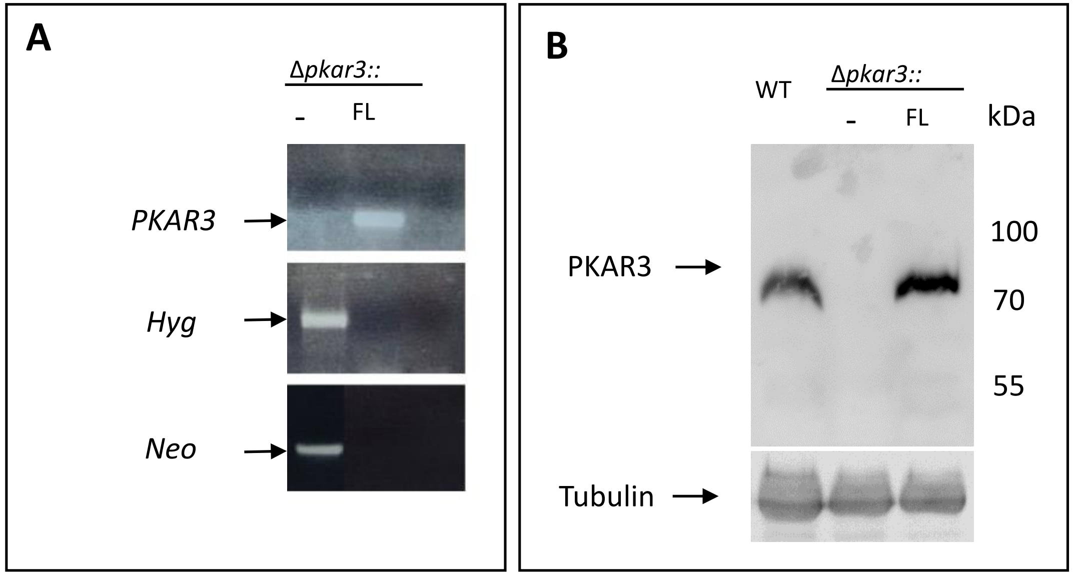
A
Δpkar3::
-
FL
PKAR3
Hyg
Neo
B
WT
Δpkar3::
-
FL
kDa
PKAR3
100
70
55
Tubulin

### Figure S3

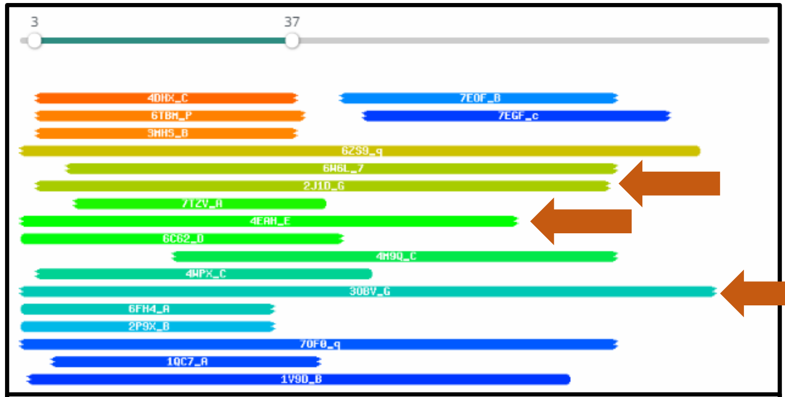


| Nr | Hit | Name | Probability | E-value | Score | SS | Aligned cols | Target Length |
|---|---|---|---|---|---|---|---|---|
| 1 | 4DHX_C | Enhancer of yellow 2 transcription factor homolog; mRNA export, TRANSPORT PROTEIN-DNA binding Protein complex; 2.1A {Hom | 91.43 | 0.52 | 31.54 | 4 | 35 | 101 |
| 2 | 6TBM_P | Transcription and mRNA export factor SUS1; Transcriptional co-activator, Histone-acetylation, TRANSCRIPTION; 20.0A {Koma | 89.13 | 1.2 | 29.61 | 4.1 | 36 | 96 |
| 3 | 3MHS_B | Protein SUS1; Multi-protein complex, Hydrolase-transcription regulator-protein binding complex, Acetylation, Cytoplasm, | 88.68 | 1.3 | 29.71 | 4.1 | 35 | 96 |
| 4 | 6ZS9_q | Growth arrest and DNA damage-inducible proteins-interacting protein 1; Mitochondria, Ribosome, mRNA, tRNA, Translation; | 82.11 | 11 | 29.7 | 7 | 90 | 198 |
| 5 | 6W6L_7 | Coiled-coil domain-containing protein 47; RIBOSOME-PROTEIN TRANSPORT complex; HET: MG; 3.84A {Homo sapiens} | 79.94 | 41 | 29.31 | 10.3 | 73 | 483 |
| 6 | 2J1D_G | DISHEVELED-ASSOCIATED ACTIVATOR OF MORPHOGENESIS 1; ACTIN ASSEMBLY, PROTEIN BINDING; 2.25A {HOMO SAPIENS} | 79.86 | 32 | 28.3 | 9.2 | 76 | 483 |
| 7 | 7TZV_A | WYL domain-containing protein; DriD, DNA repair, SOS-independent, WYL domain, wHTH, TRANSCRIPTION; 1.65A {Caulobacter vi | 71.68 | 8.5 | 25.55 | 3.3 | 34 | 198 |
| 8 | 4EAH_E | Formin-like protein 3; ATP binding, cytoskeleton, formin, FMNL3, actin, PROTEIN BINDING; HET: ATP, ACT; 3.4A {Oryctolagu | 69.79 | 29 | 27.02 | 6.3 | 66 | 402 |
| 9 | 6C62_D | AtzG; AtzE biuret hydrolase atrazine cyanuric acid Ser-cisSer-Lys hydrolase, HYDROLASE; HET: CSO; 1.95A {Pseudomonas sp. | 69.33 | 35 | 22.16 | 5.9 | 43 | 68 |
| 10 | 4M9Q_C | ARF-like GTPase; GTPase, G domain, Joubert Syndrome, cilia, HYDROLASE; HET: GNP, SO4; 2.5A {Chlamydomonas reinhardtii} | 65.59 | 38 | 21.19 | 8.3 | 59 | 227 |
| 11 | 4WPX_C | Putative uncharacterized protein; cell cycle, mRNA nuclear export; 3.31A {Chaetomium thermophilum} | 61.32 | 14 | 28.87 | 3.2 | 45 | 181 |
| 12 | 3OBV_G | Protein diaphanous homolog 1; AUTOINHIBITION, ACTIN, NUCLEATION, CYTOSKELETON, STRUCTURAL PROTEIN; HET: GLC, FRU; 2.75A | 59.26 | 110 | 24.56 | 9.7 | 90 | 457 |

Figure S4

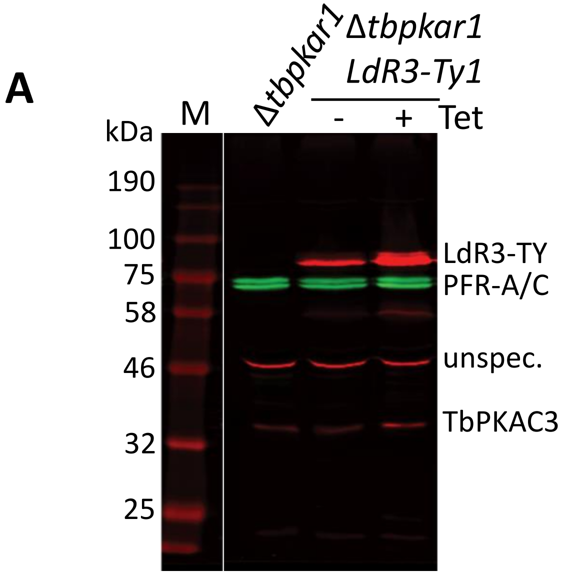
A
M
Δtbpkar1
Δtbpkar1
LdR3-Ty1
- + Tet
kDa
190
100
75
58
46
32
25
LdR3-TY
PFR-A/C
unspec.
TbPKAC3
B
WCL
sol
P
M
kDa
100
75
58
46
32
LdR3-TY
unspec.
TbPKAC3
SAXO

### Figure S5

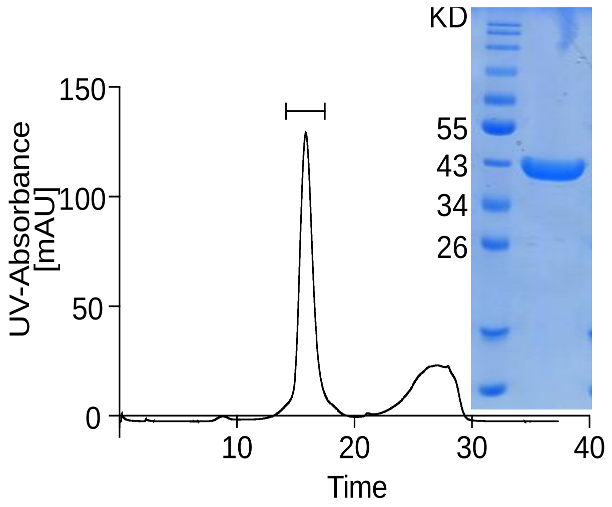

Figure S6

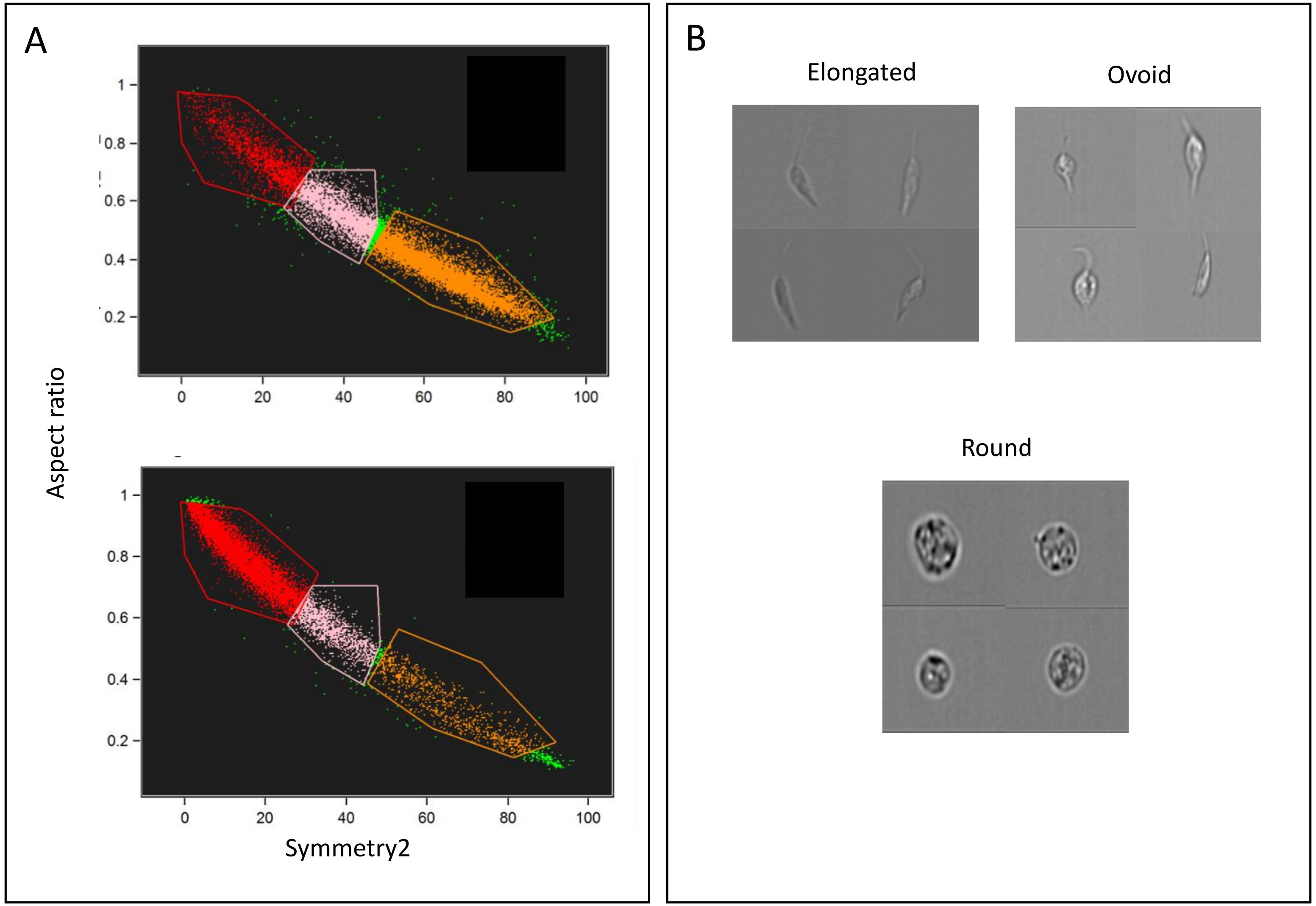
A
Aspect ratio
1
0.8
0.6
0.4
0.2
0
20
40
60
80
100
Symmetry2
B
Elongated
Ovoid
Round

Figure S7

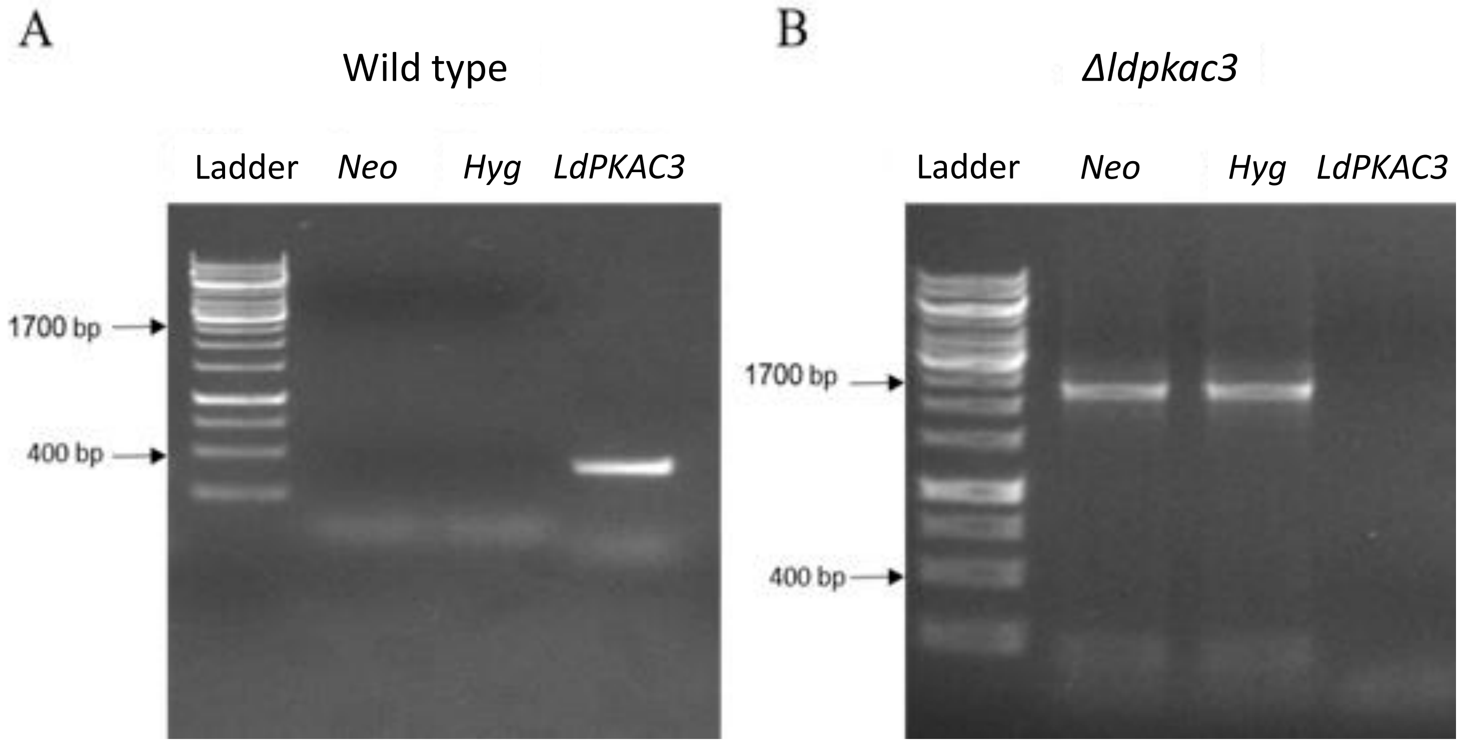
A
Wild type
Ladder
Neo
Hyg
LdPKAC3
1700 bp
400 bp
B
Δldpkac3
Ladder
Neo
Hyg
LdPKAC3
1700 bp
400 bp
